# Reward impacts visual statistical learning

**DOI:** 10.1101/2020.04.04.025668

**Authors:** Su Hyoun Park, Leeland L. Rogers, Matthew R. Johnson, Timothy J. Vickery

## Abstract

Humans automatically detect and remember regularities in the visual environment—a type of learning termed visual statistical learning (VSL). Many aspects of learning from reward resemble statistical learning in respects, yet whether and how reward learning impacts VSL is largely unexamined. In two studies, we found that reward contingencies affect VSL, with high-value associated with stronger behavioral and neural signatures of such learning than low-value images. In Experiment 1, participants learned values (high or low) of images through a trial-and-error risky choice task. Unbeknownst to them, images were paired as four types—High-High, High-Low, Low-High, and Low-Low. In subsequent recognition and reward memory tests, participants chose the more familiar of two pairs (a target and a foil) and recalled the value of images. We found better recognition when the first images of pairs have high-values, with High-High pairs showing the highest recognition rate. In Experiment 2, we provided evidence that brain responses were affected by both value and statistical contingencies. When we compared responses between the high-value first image and the low-value first image, greater activation in regions that included inferior frontal gyrus, anterior cingulate gyrus, hippocampus, among other regions were found. These findings were driven by the interaction between statistically structured information and reward—the same value contrast yielded no regions for second-image contrasts and for singletons. Our results suggested that the powerful allocation of attention in response to the high-value first image potentially enables better memory for statistically learned pairs and reward information than low-value first image.

## INTRODUCTION

Reward motivation impacts human cognition in many contexts (Haber & Knutson, 2010). Value is linked to stimuli that are critical for individuals’ survival (e.g., primary reward; water or food), but learned associations between reward and neutral stimuli can also shape one’s behavior (e.g., secondary reward; money; Daw & Doya, 2006). There is vast literature demonstrating how secondary cues, especially monetary reward, guide an individual’s cognitive processes such as memory, attention, and decision making. Higher associated value facilitates stimulus-reward memory association (Adcock et al., 2006), and features and objects that are associated with higher value capture more attention than those with low- or no rewards (e.g., Anderson, 2013; Theeuwes & Belopolsky, 2012). Individuals’ decision-making tends to optimize action so that rewards are maximized and losses minimized (Tversky & Kahneman, 1979). However, the relationship between learning and reward is typically studied in the context of learning rewarding associations, specifically, or memory of individual stimuli that are explicitly or implicitly associated to reward (Miendlarzewska et al., 2016). In the present study, we examined how learning explicitly about rewarding associations modulates the undirected and uncued learning of visual statistical associations.

Visual statistical learning (VSL) is a type of learning that reflects automatic and unsupervised extraction of statistical contingencies by the visual system (Fiser & Aslin, 2001, 2002). Prior studies suggested that humans may, in part, accomplish efficient processing of complex visual environments by learning and exploiting knowledge of visual regularities (Fiser & Aslin, 2001, 2002; Turk-Browne et al., 2005). In two early VSL studies, Fiser and Aslin (2001, 2002) found that when particular visual items co-occurred with others, subsequent recognition rates of those regularities were above chance, even though those regularities were task-irrelevant, no instructions to remember the associations were given, and the associations were not cued. A typical VSL paradigm takes place in the context of passive viewing or simple cover tasks. How VSL occurs in the context of different task demands and contexts, as it must occur in everyday life, is underexplored. Since intentional seeking and learning about rewards is so foundational to behavior, it is natural to ask how learning about reward might impact incidental learning of regularities.

Several findings support the idea of potential pathways for reward to influence VSL. Prior studies provided evidence that these two types of learning incorporate some similar associative mechanisms. For instance, both stimulus-reward and stimulus-stimulus contingencies are learned as a result of being presented for multiple times of these contingencies, intentionally or unintentionally. Further, the neural studies provide reasons to suspect reward learning and VSL may be interrelated. That is, these two types of learning have been shown to share common neural correlates, at least regionally: correlates of both reward learning and VSL have been found in hippocampus, striatum and medial temporal lobe (Aron, 2004; Delgado et al., 2000; Lansink et al., 2009; Wittmann et al., 2007). As these brain areas are known for their key roles in associative learning (Rieckmann et al., 2010), reward and visual statistical learning may share common neural substrates, in terms of extracting and binding meaningful information (or patterns) and predicting and evaluating upcoming events based on that information.

Shared neural correlates between reward learning and VSL have also been found in the lateral occipital cortex (LOC), such that greater LOC activity was shown when exposed to visual regularity vs. no regularities (Turk-Browne & Scholl, 2009) and high reward vs. low reward (Anderson, 2017). LOC is known for its role in object perception (Grill-Spector et al., 2001; James et al., 2003), but previous studies also showed greater LOC activation to attended relative to distractor or ignored objects (Vuilleumier et al., 2005; Woolgar et al., 2015). Considering prior evidence that attention modulates response patterns in higher visual areas (e.g., Murray & Wojciulik, 2004), variations in LOC as a function of both forms of learning may be related to attentional processing during object perception (see also Stokes et al., 2009). Neural activity in LOC in response to both types of learning may imply one potential pathway of reward to influence on VSL, as attention may play an important role in their relationship. Indeed, the role of attention in each of these phenomena is compelling enough to believe that reward may impact VSL on the basis of how reward shapes attention, alone.

According to previous studies, selective attention plays an important role in VSL, as selective attention to stimuli is required for VSL to occur (Baker et al., 2004; Turk-Browne et al., 2005). Baker et al. (2004) found that visual regularities were not learned in the absence of selective attention, such that when the memory of target-distractor combinations was tested, no learning was observed when participants paid attention to only one target location, but VSL occurred when participants paid attention to both of target and distractor locations. Turk-Browne et al. (2005) also found the learning of regularities occurred only with an attended color stream when participants were exposed to an interleaved stream that composed of attended-and unattended-color (but see Musz et al., 2015). Based on prior studies, we assume learning of structural information does not occur in a uniform way. Rather, selective attention may be required to process such information, with the degree of selective attention determining the strength of learning.

Mounting evidence suggests that rewards bias attention towards stimuli that have been associated with those rewards as well (e.g., Anderson et al., 2013; Won & Leber, 2016). High value associations weakened the effect of the attentional blink, meaning that high reward biases one’s attention (Raymond & O’Brien, 2009), and an oculomotor capture of stimulus-reward associations revealed that a stimulus associated with a high amount of monetary reward captures more attention than that associated with a low reward (Theeuwes & Belopolsky, 2012). Anderson et al. (2013) found greater attention was driven to items that were previously learned as high value items than low value items even though value information is no longer relevant to the task, and participants were not able to explicitly remember the association between stimulus and reward outcomes.

Based on previous research, we predicted that when different amounts of reward are embedded in visual regularity, the rewards may interact with VSL, and selective attention potentially plays an important role in mediating this interaction. Reward is a powerful influence on the allocation of attention (Theeuwes & Belopolsky, 2012), and VSL is dependent upon selective attention to the constituent items (Turk-Browne et al., 2005). Therefore, VSL may be modulated by reward information, with high reward items driving more attention than low reward items and resulting in stronger memory formation of the constituent items that are associated with high reward.

We further predicted that the effect of reward might be especially potent when the high reward item is in the first position in a temporally presented pair sequence. In VSL, the position of an item in a stereotyped sequence seems to determine the neural response profile to that item (Turk-Browne et al., 2010). Turk-Browne et al. (2010) showed that the right anterior hippocampus and medial temporal lobe showed enhanced responses when the first picture of a pair appeared (i.e. predicting the stimuli) as compared to novel singletons. These results suggest that during the acquisition of statistical regularities, the first item of the structured information plays an important role in predicting and evaluating subsequent items.

Hence, when reward is embedded in VSL sequences, reward may evoke different responses according to the position of the structured information it is associated with. In particular, higher reward that is specifically associated with early items in a temporal sequence may aid visual statistical learning. If attentional processing is involved in this interaction, greater activations may be found in brain regions in frontal and parietal areas, such as inferior frontal gyrus, precentral gyrus, and anterior cingulate gyrus, that are known for their roles in attentional network and cognitive control (e.g., Corbetta & Shulman, 2002; Fockert et al., 2004; Kan & Thompson-Schill, 2004), in addition to the LOC.

To our knowledge, Rogers et al. (2016) is the only work to examine the relationship between monetary reward and VSL directly. Despite finding evidence of visual statistical learning, the amount of reward associated with stimuli and sequences did not affect the strength of VSL in their studies, suggesting that reward processing and VSL were operating independently. However, the manipulation of reward, in that case, may have been too subtle for participants to process reward contingencies in a VSL paradigm. Therefore, to motivate learning and enhance participants’ performance, we employed a risky choice task (e.g., Clark et al., 2009), which is more likely to lead to in-depth processing of reward information. With this manipulation, we expected that participants would be more engaged in the task, and enhanced learning would be observed for both reward and statistical information.

In the present study, we examined how reward modulates VSL. To clearly see the interaction between varying rewards (i.e., high vs. low) and the position of an item in a structured sequence, we used pairs presented in temporal succession to instantiate statistical regularities, but pairs were constructed with different reward variations (i.e., High-High, High-Low, Low-High, and Low-Low). We found higher recognition rates for pairs when the first image of a pair had a high-value, which suggested the high value of the first item in a pair enhances learning (or low-reward impairs learning). The neural correlates linking reward variations to statistical regularities was examined in Experiment 2. Using event-related fMRI, we measured brain responses to images that were associated with both varying levels of reward (i.e., high vs. low) and sequential contingencies (i.e., the first or second image in a pair, or singletons). The first image with a high-value, in comparison to the first image with a low-value, led to greater activity in areas including IFG, left ACC, LOC, OFC, accumbens, hippocampus, and putamen. This serves as circumstantial evidence that reward may play a role similar to selective attention in VSL, or it may affect VSL by shaping selective attention, since many of these neural correlates are shared with those evoked by manipulations of attention. Importantly, we also provided evidence that the differences between the high-value first image and the low-value first image are not driven solely by the value difference, but by an interaction of predictiveness and value.

## EXPERIMENT 1

The aim of Experiment 1 was to examine the influence of learned value on VSL by embedding different amounts of reward into structured pairs (i.e., High-High, High-Low, Low-High, and Low-Low reward pairs) that always co-occurred temporally in a sequence of decisions. After participants learned the value in a temporally structured sequence, we tested recognition for each type of pair, allowing us to examine how the high- or low-reward association might interact with the location of reward (i.e., first or second) in structured pairs.

### Method

#### Participants

All procedures were approved by the University of Delaware Institutional Review Board. Thirty-three University of Delaware students who were 18-40 years of age participated for course credit or cash. At the last phase of Experiment 1, participants’ memory for the image value was measured. Pilot data suggested that reward memory recognition judged by the third phase was almost always above chance levels. Since it was crucial for participants to have a memory of reward associated with constituent items to judge reward effects on VSL, we established exclusion criteria based on last-phase performance. Two participants were excluded because they did not show above chance (50%) reward memory recognition rate.

#### Stimuli and Apparatus

Experiment 1 was run on Windows 10 with a 24-inch LCD monitor with a resolution of 1920 × 1080. The experiment was programmed in MATLAB with Psychophysics Toolbox v. 3 (Brainard, 1997; Kleiner et al., 2007). We used 32 fractal images as novel visual stimuli. Images were randomly assigned into structured sequences (i.e. pairs) between participants. Stimuli were 200 pixels x 200 pixels, and participants sat approximately 57 cm from the monitor (images subtended approximately 5° of visual angle).

#### Procedure

Experiment 1 consisted of three phases. Participants performed 1) a learning phase followed by 2) a surprise pair recognition phase. In the last phase, they completed 3) a reward memory test, which asked participants to explicitly recall the value of each image (i.e., high or low; two-alternative forced-choice task). Before the experiment began, participants were given instructions about the learning task. However, no information was provided to participants about the subsequent memory-test phases prior to completing the learning phase. Regarding incentives, participants were told that during the learning phase, points would be shown on the screen based on their choice (the description is below). Points added up over time and they would get money based on their point totals. At the end of the experiment, the points were converted to maximum of $10. Total points (maximum of 3200) were divided by 320 to derive this value. Participants were informed that points would be converted to money at the end of the experiment (up to $10 total), but not of the exact conversion rate.

During the learning phase (Fig 1A), images were presented at the center of the screen, sequentially. Participants were instructed to do a risky choice task, in which they learned the values (high or low) of fractal images through trial-and-error. For each image, participants needed to make a choice (phrased as a “gamble”) of “Yes” or “No.” If they chose “Yes” (press the Z button on the keyboard), they had a 50% chance of winning nothing (0 points) and a 50% chance of winning points. Importantly, “high-reward” images were associated with a 50% chance to win 10 points, while “low-reward” images were associated with a 50% chance to win two points. If they chose “No” (pressed the M button on the keyboard), they always got one point and, importantly, were able to see what they could have gained (i.e., 0, 2, or 10) if they chose “Yes” on that trial. This way, they were still able to learn 1) the associated value (if two or 10 points were assigned on that trial) and 2) whether they won by not choosing “Yes” on that trial (if 0 was assigned on that trial). If they could not choose within two seconds, it was counted as “Miss.”

**Figure 1.**
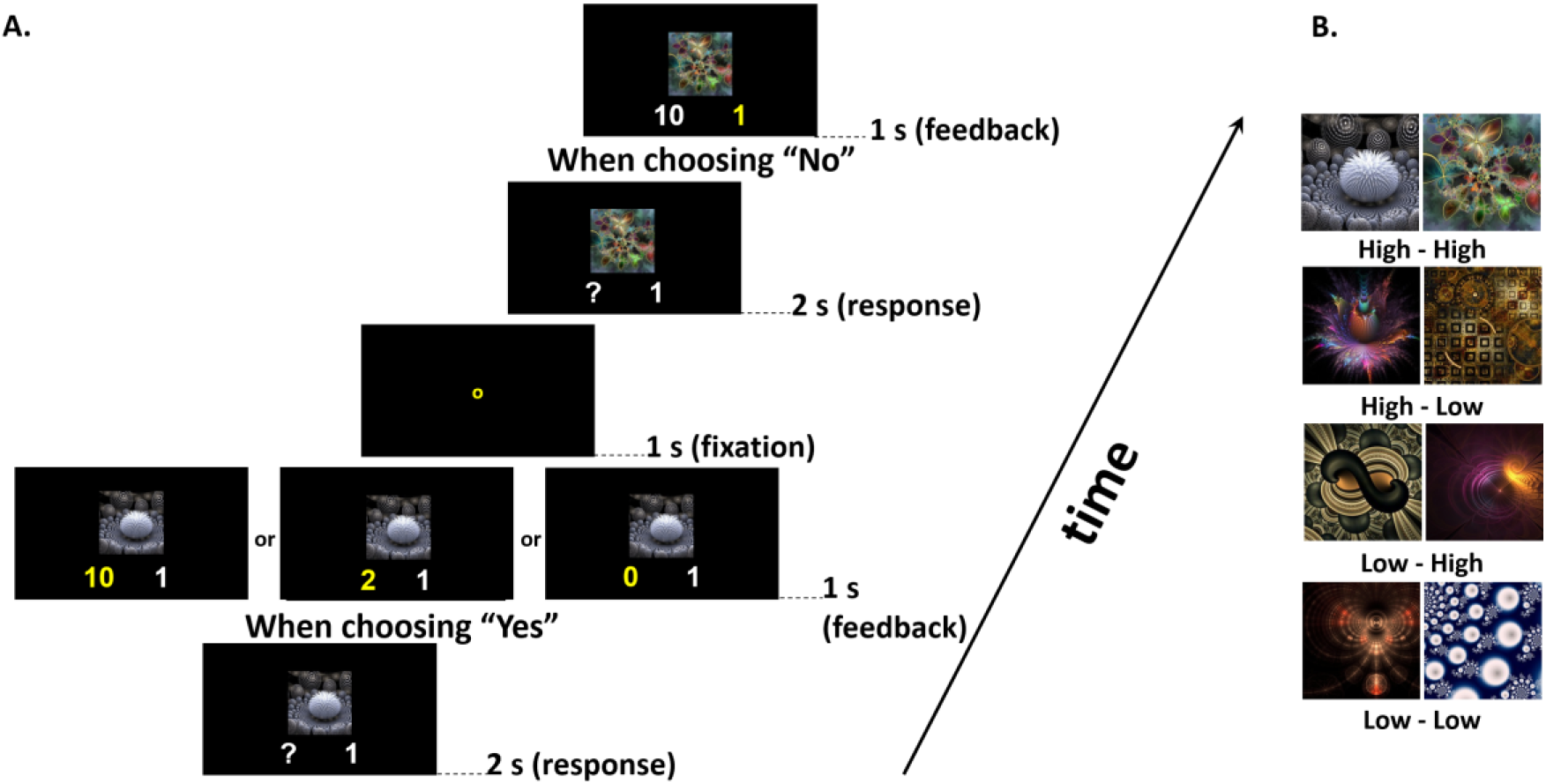
A) General procedure of the learning phase in Experiment 1. B) Pairs were equally divided into four reward variations (High-High, High-Low, Low-High, and Low-Low).

Unbeknownst to participants, we paired images so that some images always predicted other images on the following trial. This led to four types of pairings (High-High, High-Low, Low-High, and Low-Low; Fig. 1B). All structured pairs were pseudo-randomized within the stream such that no immediate repetition of a pair (e.g., ABAB) or two sets of pairs (e.g., ABEFABEF) could occur. The 32 fractal images (16 pairs) were repeated four times within each block. With a total of five blocks, each image/pair appeared a total of 20 times. The 16 pairs were equally divided into four of each of the pairing conditions.

Following the learning phase, the recognition phase began. Participants were given on-screen instructions before they began the recognition phase. This phase involved a two-alternative forced-choice task in which participants were asked to choose which of two two-image sequences was more familiar (Fig. 2A). One of the sequences was a sequence of a target pair, and the other one was a sequence of a foil pair. The target pair was a structured pair that was presented multiple times during the learning phase (e.g., AB, CD, EF, etc.). Foil pairs were recombined from pairs constructed from using the first image of one target pair and the second image of another target pair (e.g., AD). Each target and foil pair were presented four times during the test phase. We constrained each target pair type (in terms of reward) to match with all types of foil pairs (e.g., High-High (target) vs. High-High (foil); High-Low (foil); Low-High (foil); Low-Low(foil)) in each presentation. No feedback was given during this phase, and participants had unlimited time to respond.

**Figure 2.**
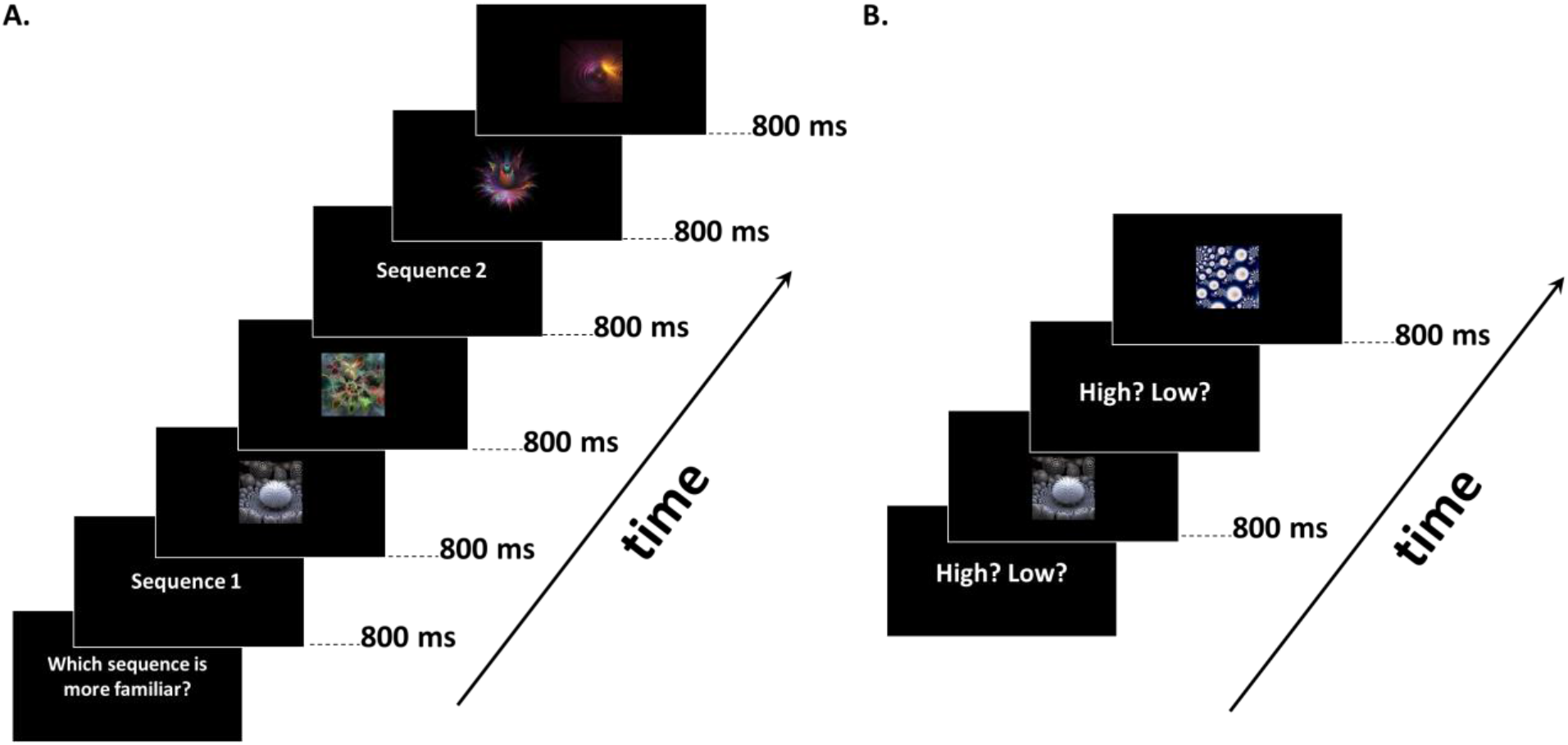
General procedure of the memory tests. (A) Example of the recognition test. (B) Example of the reward memory test.

After the recognition phase, participants were asked to remember the value of all images that they saw during the learning phase and choose whether they had high or low-values in a two-alternative forced-choice paradigm. All 32 images were presented one by one in a random order (Fig. 2B), with no time constraints and no feedback provided.

## RESULTS

A two-way repeated measures ANOVA (value of image × block) on risky choice proportion (i.e., choosing yes) showed a significant main effect of value of image, F (1, 30) = 48.04, p < .001, ηp^2^ = .616 (but no main effect of blocks, F <1) and an interaction between block and value, F (4, 120) = 12.48, p < .001, ηp^2^ =.3. Proportion of making a risky choice to high-value images gradually increased across blocks, and the opposite was observed with low-value images (Fig 3).

**Figure 3.**
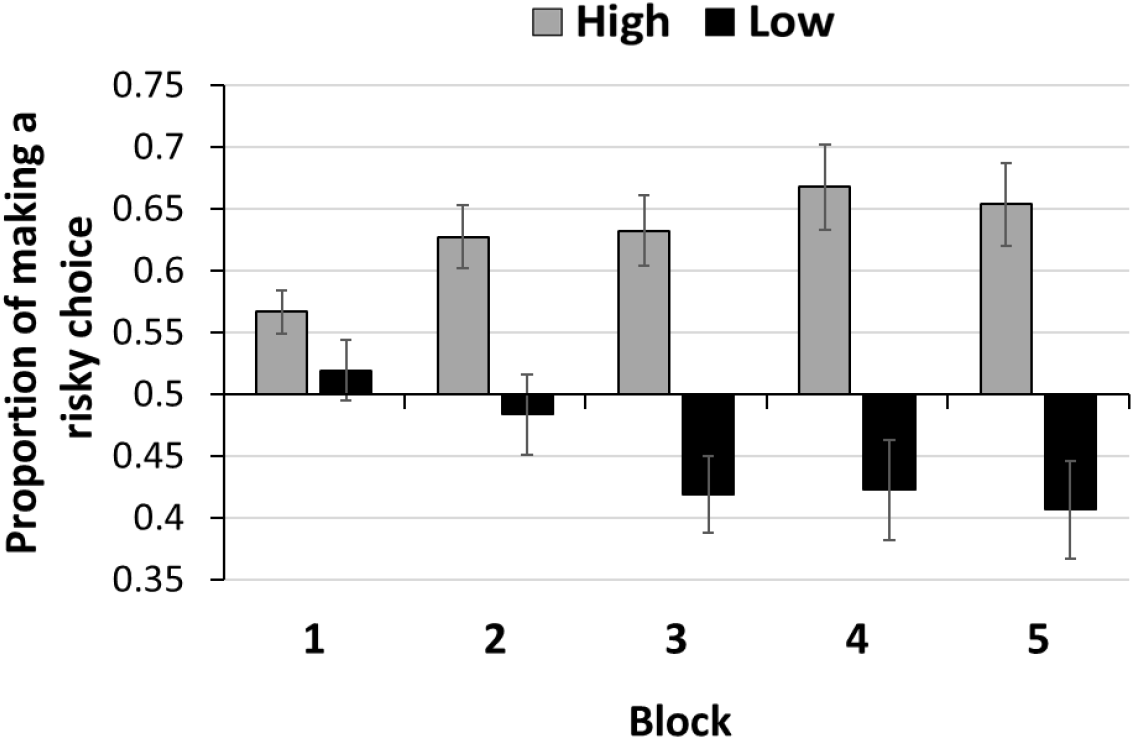
Proportion of making a risky choice throughout blocks. In this and all other figures, error bars represent standard error of the mean.

In regards to the recognition phase, a one-sample t-test against chance (50%) yielded significant learning only for the High-High condition, t(30)= 2.71, p =.01, d=.49. In addition, with a 2 (value of first image, high or low) × 2 (value of second image, high or low) repeated measures ANOVA, we only found a significant main effect of the first image such that there was better recognition when the first image of a pair was a “High” image, F(1, 30) = 6.41, p = .017, ηp^2^ = .17 (Fig. 4). No main effect of the second image nor interaction were found (F < 1). Bayesian Repeated Measures ANOVA (using a position of image as a factor; the first or the second position in a pair) showed evidence that the effect of value of the second image favored the null hypothesis (main effect of the second position, BF01 = 3.425; interaction, BF01 = 2.725), indicating mild evidence against the possibility of an effect of second-image value. To ensure that results were not impacted by foil pair value, we conducted a 2 (value of first image, high or low) × 2 (value of second image, high or low) repeated measures ANOVA based on foil type, which resulted in no significant main effects and no interaction of foil value on recognition accuracy, all F <1.

**Figure 4.**
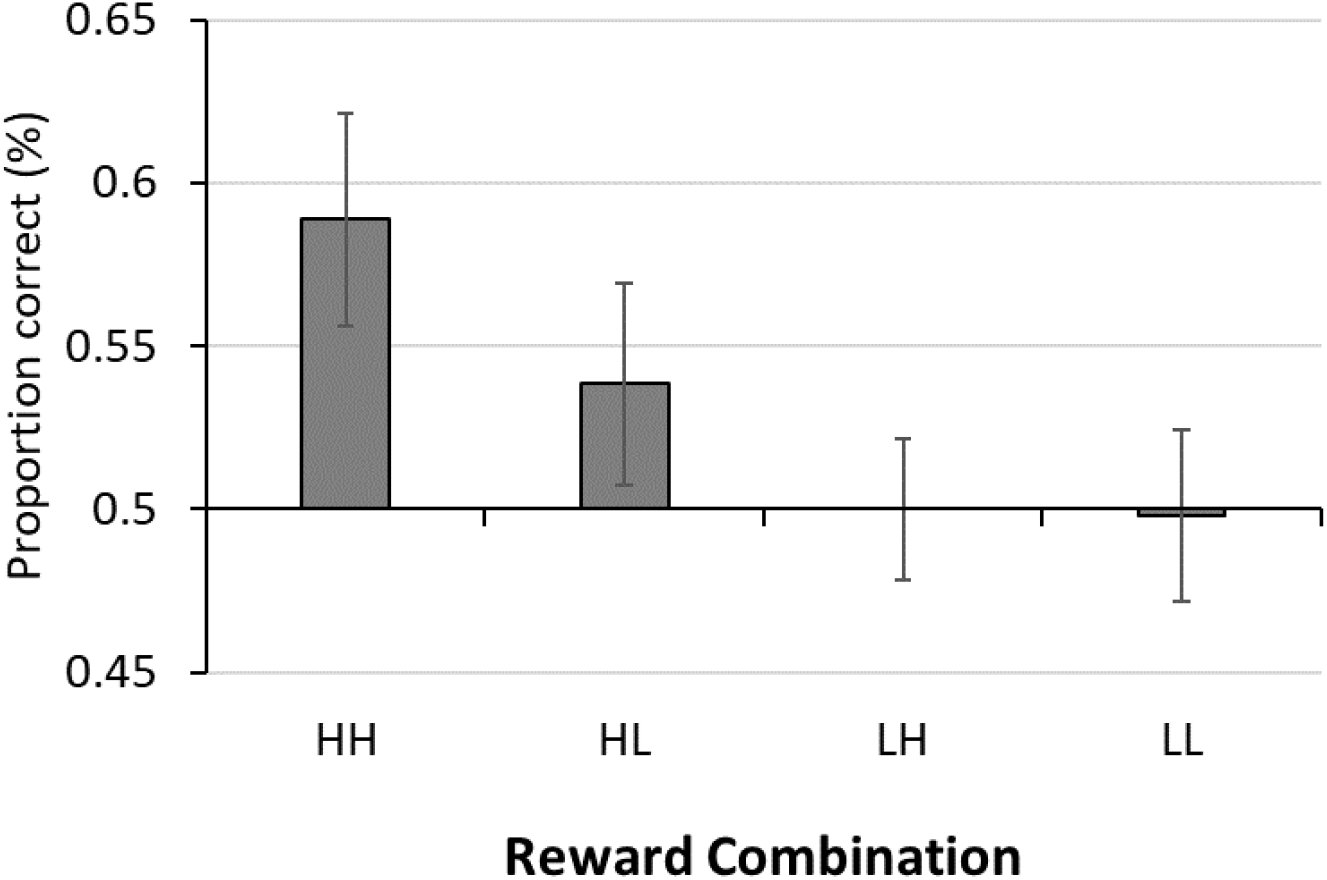
Accuracy at choosing target pairs over foil pair in four reward variations.

## DISCUSSION

Previous research found no differences in VSL amongst no-, low-, or high-reward conditions (Rogers et al., 2016). However, previous efforts did not explicitly draw attention to value during exposure to statistical associations. In the current study, using a risky choice task, the participants’ task was to learn the value of images, which drew attention explicitly to reward during exposure. Under these constraints, we found better recognition for pairs when the first image of the pair was a high-reward image.

A number of mechanisms might explain this finding, with variations in attention caused by associated value being one candidate. In VSL, the first item of structured pairs plays an important role in predicting and evaluating subsequent outcomes during the acquisition of statistical regularities (Turk-Browne et al., 2010). Reward could impact VSL by drawing intense attention to the high-reward image that was located in the first position of a pair. As we do not see any benefit for pairs where the high-reward image appeared second (i.e., Low-High pairs), we speculated that value information might interact with VSL because attention is engaged with greater frequency and/or intensity when the first image of a pair is associated with high-reward, in advance of the predictable second image. This in turn enables learning of the association. On the other hand, if the first image of a pair did not receive such priority (i.e., the low value first image), VSL may not be fully engaged. In Experiment 2, we investigated the neural correlates of how reward variations affect the learning of statistical regularities and probe the underlying mechanisms of our finding that reward associations shape VSL. We mainly focused on how attentional processing is involved in this interaction and sought to find whether high reward would affect VSL by shaping the propensity of the system to learn contingencies.

## EXPERIMENT 2

To investigate the neural basis of how reward impacts VSL, we measured brain responses to visual images that were associated with both varying levels of reward and sequential contingencies, using event-related fMRI. We examined the neural activation of the first and the second image in pairs, and how it differed according to the amount of reward (high vs. low). We also compared images with structural information (i.e., pairs) and without such information (i.e., singletons) in each of high and low-value (e.g., high paired images vs. high singleton; low paired images vs. low singleton), and asked how the varying level of reward affected the processing of statistically structured information.

## METHOD

### Participants

Thirty University of Delaware students who were 18-40 years of age each participated in one 2-hour long experimental session (mean age: 21.6; 22 females). One participant did not show above chance levels of learning in the last reward memory phase, so that participant was excluded from further analysis. All participants were right-handed, reported having normal color vision, and were compensated $20/hour. All procedures were approved by the University of Delaware Institutional Review Board.

### Stimuli, Apparatus, and Procedure

In Experiment 2, a total of 48 fractal images were used. 32 images were assigned to 16 pairs, and the remaining 16 images were used as singletons. The added singletons allowed us to directly compare the differences in neural activity for images that contained statistical structure information and images that do not.

There were four phases, 1) the risky choice task (i.e., the learning phase), 2) the passive viewing task, 3) the recognition test, and 4) the reward memory test. Participants performed the risky choice task and the passive viewing task inside of the scanner, and two memory tests were performed outside of the scanner. The rules of the risky choice task were identical to Experiment 1. However, the procedure timing was modified to accommodate fMRI analysis. In Experiment 2, there were four runs of the risky choice task, and in each run, a new set of four pairs and four singletons were presented, with each repeating six times within the block. We chose six repetitions based on prior studies showing evidence of learning even with a small number of repetitions (Turk-Browne et al., 2010), and so that we could introduce new images in each run. Additionally, we included jittered intervals between 1) the choice phase and feedback phase of the trial and 2) the feedback phase of the trial and the next image presentation (Fig. 5). Jittered intervals consisted of 2s, 3s, 4s, or 5s, and they were evenly divided across conditions and presented in a randomized order. During the risky choice task phase, participants responded with an MRI-compatible button box. Following the four learning runs, a passive viewing run was performed. In this run, all 48 images were presented one more time, with each of the 16 pairs presented in pair-wise order and 16 singletons randomly presented in between pairs. Participants were asked to focus on each image but otherwise passively view them. Each image was presented for one second followed by a jittered interval [2s, 3s, 4s, or 5s] ^1^. Despite the fact that these modifications (e.g., newly presented pairs and singletons in each run, jittered intervals, and passive viewing task) may yield different patterns of behavior results as compare to Experiment 1, we modified the experimental settings to follow the best approach to measure the neural responses of how rewards impacts VSL.

**Figure 5.**
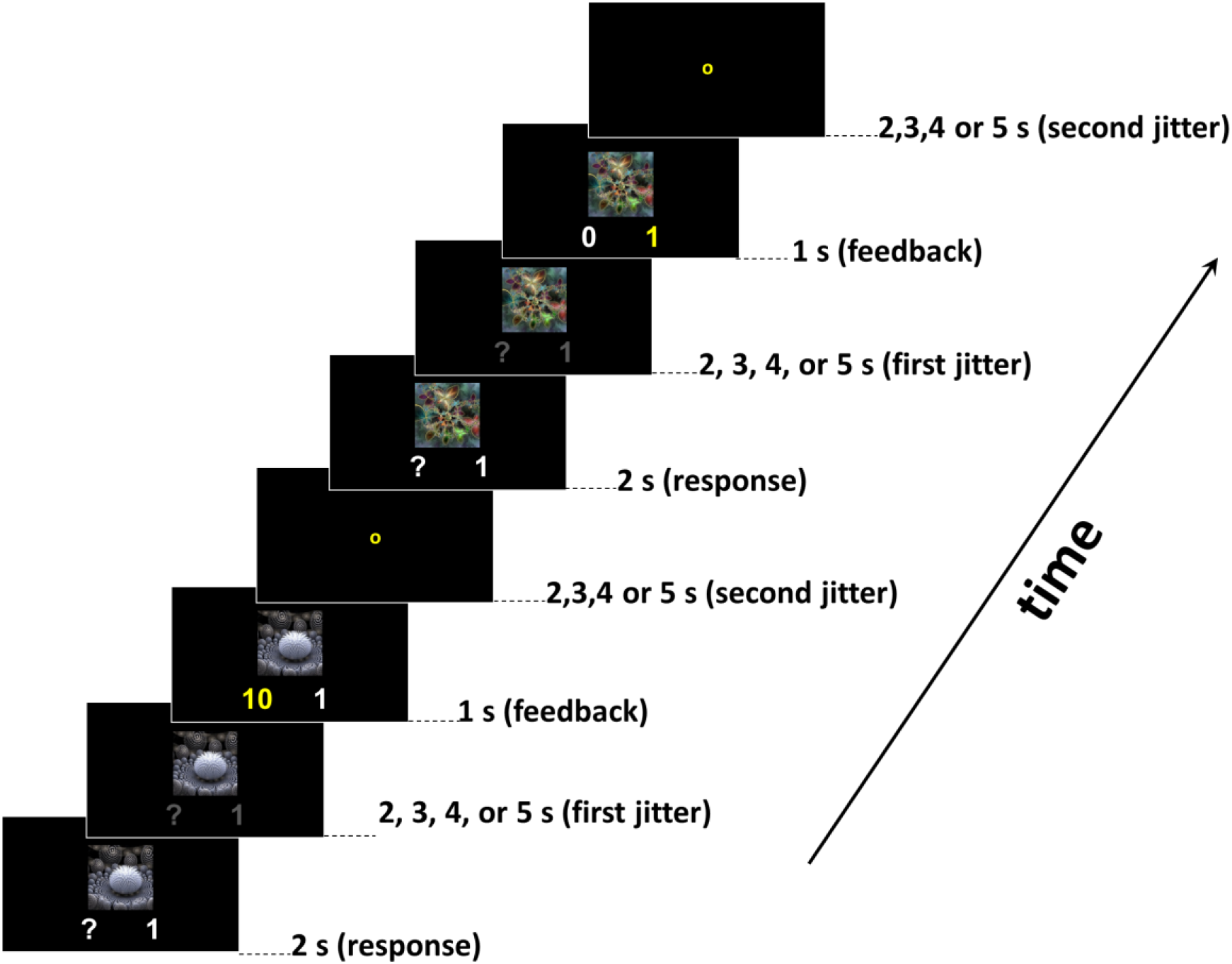
General procedure of the learning phase in Experiment 2.

After all runs, participants completed the recognition test and reward memory test outside of the scanner. The procedures for the recognition and reward memory tests were the same as Experiment 1, and all 48 images (including singletons) were shown in the reward memory test. With the experimental design modifications described above, we anticipated the possibility that the recognition and the reward memory tests may not result in the same pattern. The incentive was provided based on the points participants earned during the risky choice task, and points were converted to a maximum of $15 (i.e., total points, maximum of 900, were divided by 60 to determine payout). Participants were informed at the beginning of the experiment of the possible reward and that points would be converted to cash rewards.

### Data acquisition

Neuroimaging data were acquired on a 3T Siemens Prisma system using a 64-channel head/neck coil. One high-resolution T1-weighted MPRAGE structural image was collected (0.7 mm isotropic voxels) for anatomical information. Functional scans consisted of a T2*-weighted Siemens Multiband (multiband factor of 8) EPI sequence with 80 slices acquired in an interleaved manner, and with an oblique axial orientation (approximately 25° from anterior commissure/posterior commissure line). The in-plane resolution was 2.0 mm × 2.0 mm, and slice thickness was 2.0 mm with no skip (TR=1 s, TE = 32 ms, flip angle 61°), resulting in isotropic voxels. Each learning run consisted of 784 volumes and lasted 13 minutes and four seconds, and each passive viewing run contained 237 volumes and lasted three minutes and 57 seconds.

### Structural and Functioning Processing

Data analyses were performed using fMRIB Software Library (FSL, www.fmrib.ox.ac.uk/fsl) version 5.0.9, FMRI Expert Analysis Tool (FEAT) version 6.0 (Jenkinson et al., 2012), and the AFNI software package (Cox, 1996). For structural scans, we first performed skull-stripping by using BET (Smith, 2002), and then registered to a standard MNI152 2-mm template. For functional runs, data were first de-obliqued (AFNI’s 3dWarp) and re-oriented to match the standard template (fslreorient2std). Then, data were motion corrected, smoothed (8 mm FWHM Gaussian kernel), and high-pass temporal filtered with a 100s cutoff.

At the first-level analysis of the risky choice task phase, a total of 116 runs (four runs, 29 participants) were modeled using a standard GLM approach. Fifteen explanatory variables (EVs) were set up: HH-First, HH-Second, HL-First, HL-Second, LH-First, LH-Second, LL-First, LL-Second, High-Singleton, Low-Singleton, Choice-Yes-Win, Choice-Yes-Lose, Choice-No-Win, Choice-No-Lose, and the first presentation of each image as a regressor of no interest. The first presentation of all images was not included in the reward/location variables, because there had been no opportunity to learn either associated value or statistical contingency. For the passive viewing task, a total of 45 runs (one run: 21 participants; three runs: eight participants) were modeled using a standard GLM. Ten explanatory variables (EVs) were set up: HH-First, HH-Second, HL-First, HL-Second, LH-First, LH-Second, LL-First, LL-Second, High-Singleton, Low-Singleton. Regressors were unit-height boxcar functions that modeled the appearance of image (two seconds duration) or the response / outcome (two seconds duration), and were convolved with a double-gamma canonical hemodynamic response function. A second-level, fixed-effect analysis was then used to combine across four learning runs within each participant for the learning phase and up to three passive viewing runs. Finally, a third-level mixed-effects analysis was used to combine participants’ data. Third-level results were cluster-corrected for multiple comparisons using Randomise, FSL's nonparametric permutation testing tool (Jenkinson et al., 2012), with 5000 permutations and threshold free cluster enhancement (TFCE). Results are FWE-corrected within each analysis.

Our primary interest was examining any effect uniquely driven by the high-value first images (H1) compared to the low-value first images (L1), to uncover activity putatively associated with attention-guided or prioritized processing coinciding with reward and order. We also ran contrasts to investigate any differences between high/low-value images that appeared with or without statistical structure (e.g., H1 or H2 > High-value singleton (Hsin); L1 or L2 > Low-value singleton (Lsin), and vice versa). This approach allowed us to explore the potential for reward to influence statistically structured or unstructured images (i.e., pairs vs singletons), as the additional associative information bound to structured images (or lack thereof for singletons) may predict learning based on their learned status as a high or low reward image. For the passive viewing phase, we focused on whether there is any relationship between reward contingencies and serial position even when the risky choice task was removed. If so, it would suggest that reward-associated structured or unstructured images continue to be represented uniquely outside of reward-related contexts.

## RESULTS

### fMRI data

#### Learning phase

To explore the potential impact of reward on early constituent item in a temporal sequence, we first uncover any differences in neural responses between the high-value first image (H1) and the low-value first image (L1) in pairs, to examine whether differences would be consistent with differences in attentional engagement. Secondly, we were interested in contrasting any such observations with differences that might arise in response to high-value second images (H2) vs. low-value second images (L2), and high-value singletons (Hsin) vs. low-value singletons (Lsin), to ask whether structure modulated this response.

The contrast of the high-value first images versus the low-value first images (i.e., H1 > L1) yielded significant clusters in middle temporal gyrus, superior temporal gyrus, parahippocampal gyrus, temporal fusiform cortex, hippocampus, amygdala, thalamus, orbitofrontal cortex (OFC) (all bilaterally) as well as right inferior frontal gyrus (IFG), left lateral occipital cortex (LOC), right accumbens, right putamen, left anterior cingulate gyrus (ACC), and left paracingulate gyrus (Table 1 and Figure 6). To examine whether these results were driven solely by the value difference (i.e., high vs. low), we contrasted the activity provoked by the high-value second images with that in response to the low-value second images (i.e., H2>L2 and L2>H2), but no significant difference was observed. There was also no significant difference between high-value singletons and low-value singletons (i.e., Hsin>Lsin and Lsin>Hsin). Additionally, a statistical comparison of the interactions between 1) (H1-Hsin) and (L1-Lsin), and 2) (H1-L1) and (H2-L2) was derived. The contrast of (H1-Hsin) > (L1-Lsin) yielded greater activation in the right postcentral gyrus, right precentral gyrus, left middle temporal gyrus, right superior temporal gyrus, left hippocampus, left amygdala, and other regions (Table 2 and Figure 7). The lack of any observable difference between high-value singleton and low-value singleton, and the significant interaction in many regions, supports the conclusion that H1 > L1 outcomes are not driven solely by the value difference, but rather an interaction between statistical regularity and value differences. With the contrasts of (H1-L1) and (H2-L2), no significant clusters were observed when using Randomise^2^. The contrast of (H2 – Hsin) and (L2 – Lsin) revealed no significant clusters, which again supports the idea that the value difference is not the only factor that drives the findings from H1 > L1.

**Table 1.** Result of the contrast with H1>L1. In this and all other tables, clusters with five or fewer voxels were not reported.

| **Anatomical Label** | **Hemisphere** | **Cluster size**<br>**(voxel)** | **P value**<br>**(TFCE)** | **Peak MNI,**<br>**mm** |
| --- | --- | --- | --- | --- |
| **High value first image (H1) > Low value first image (L1)** |  |  |  |  |
| Middle Temporal Gyrus, Superior<br>Temporal Gyrus, Parahippocampal<br>Gyrus, Temporal Fusiform Cortex,<br>Hippocampus, Amygdala,<br>Thalamus, Orbitofrontal Cortex | Left | 9220 | 0.009 | -54, -28, -4 |
| Parahippocampal Gyrus, Temporal<br>Fusiform Cortex,<br>Hippocampus, Amygdala,<br>Thalamus, Accumbens | Right | 2360 | 0.014 | 26, -34, -16 |
| Middle Temporal Gyrus, Superior<br>Temporal Gyrus,<br>Supramarginal Gyrus, Planum<br>Temporale, Parietal Operculum | Right | 890 | 0.034 | 72, -32, 2 |
| Orbitofrontal Cortex, Inferior<br>Frontal Gyrus | Right | 122 | 0.041 | 28, 20, -22 |
| Frontal Pole | Left | 54 | 0.04 | -16, 50, 42 |
| Cingulate Gyrus | Left | 23 | 0.041 | -2, -12, 34 |
| Occipital Fusiform Gyrus | Right | 13 | 0.046 | 30, -68, 0 |
| Lateral Occipital Cortex | Left | 8 | 0.048 | -28, -88, 20 |
| Paracingulate Gyrus | Left | 6 | 0.05 | -4, 46, 8 |

**Figure 6.**
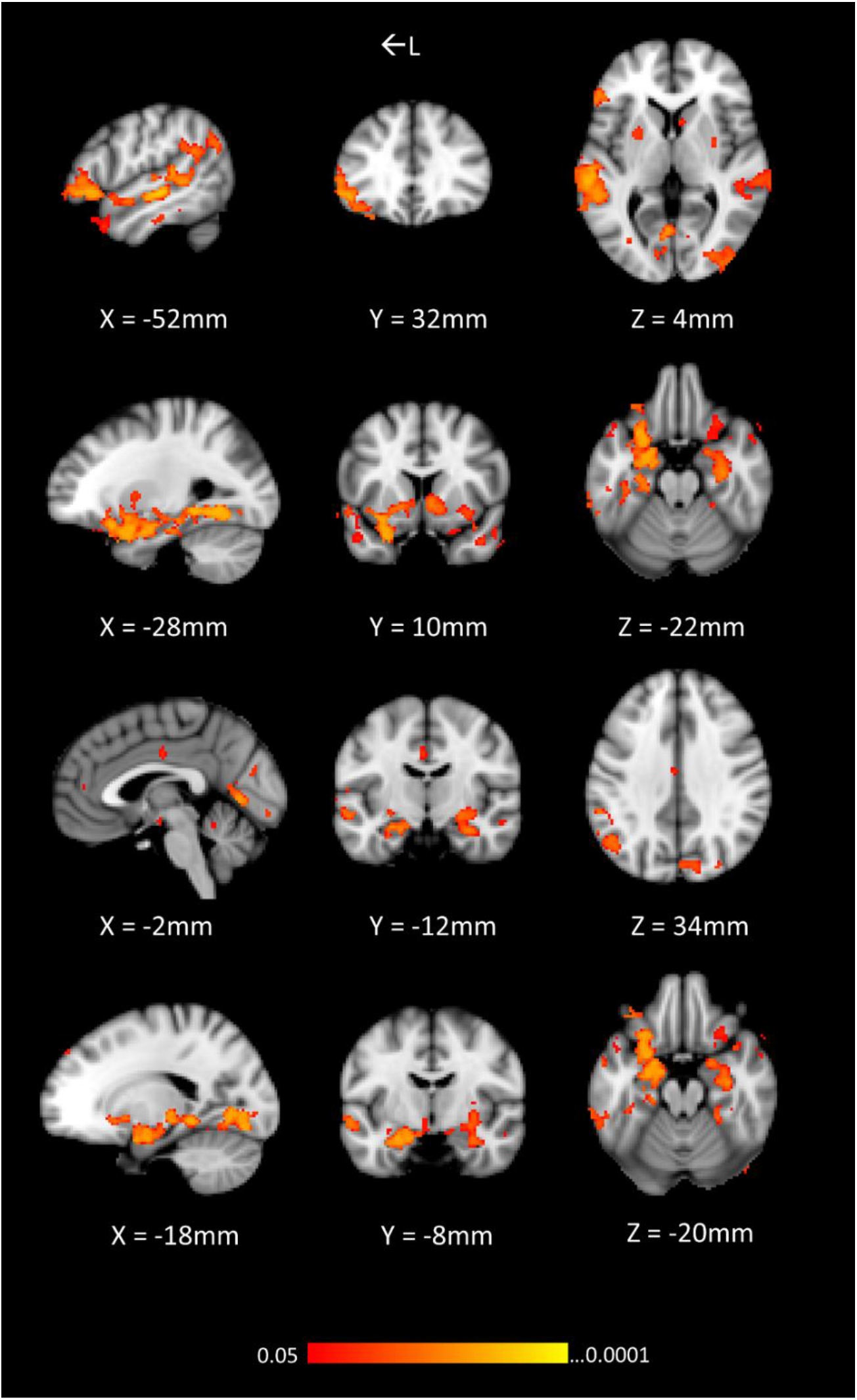
The H1 > L1 contrast yielded clusters that included middle temporal gyrus, superior temporal gyrus, parahippocampal gyrus, temporal fusiform cortex, hippocampus, amygdala, thalamus, OFC (all bilaterally) as well as right IFG, right putamen, left ACC, left LOC, right accumbens, and left paracingulate gyrus. From top to bottom, coordinates are centered on left IFG, left OFC, left ACC, and left hippocampus. In this and all other figures, coordinates are in MNI standard space.

**Table 2.**
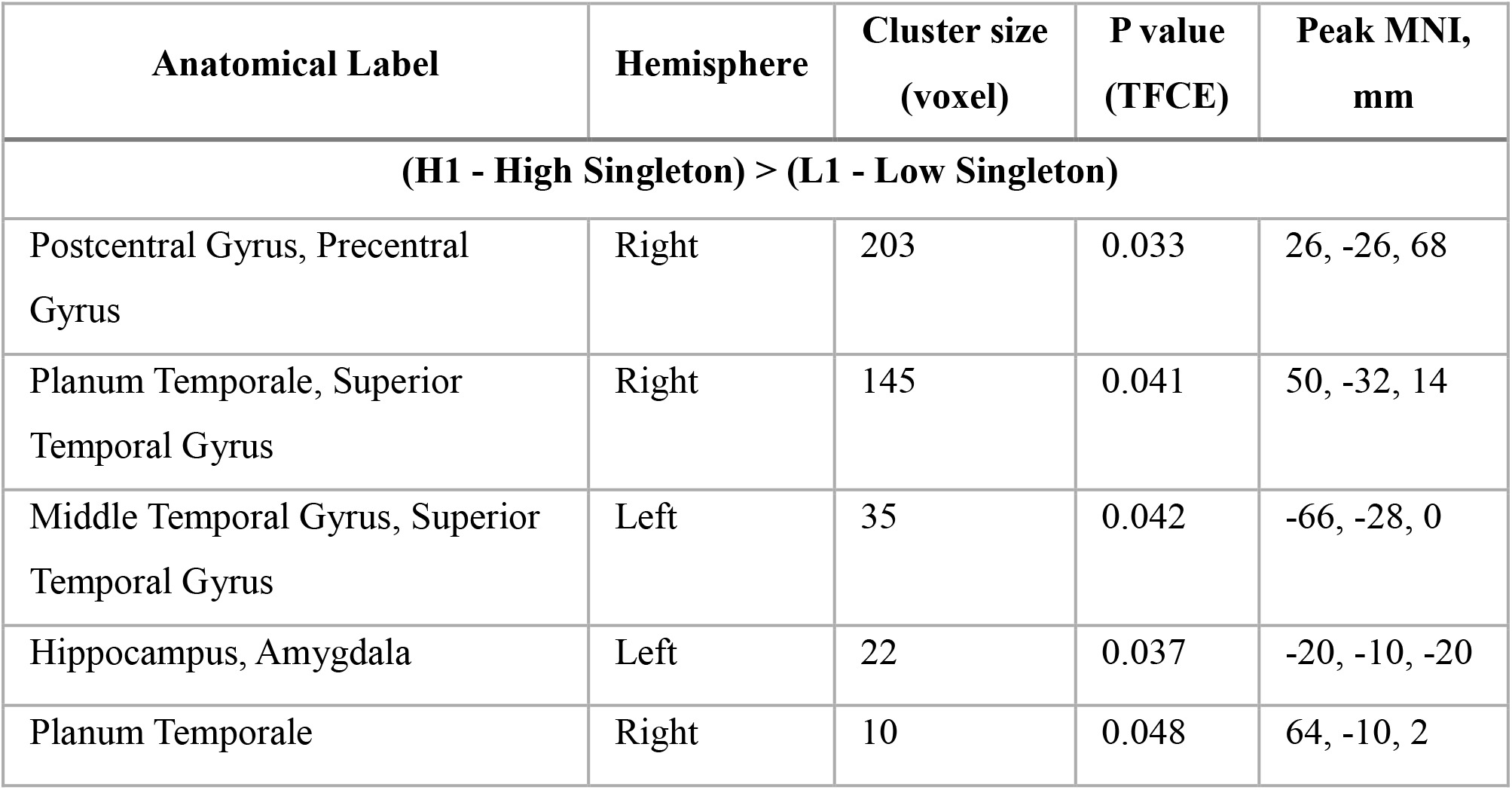
Result of the contrast with (H1-Hsin) > (L1-Lsin).

**Figure 7.**
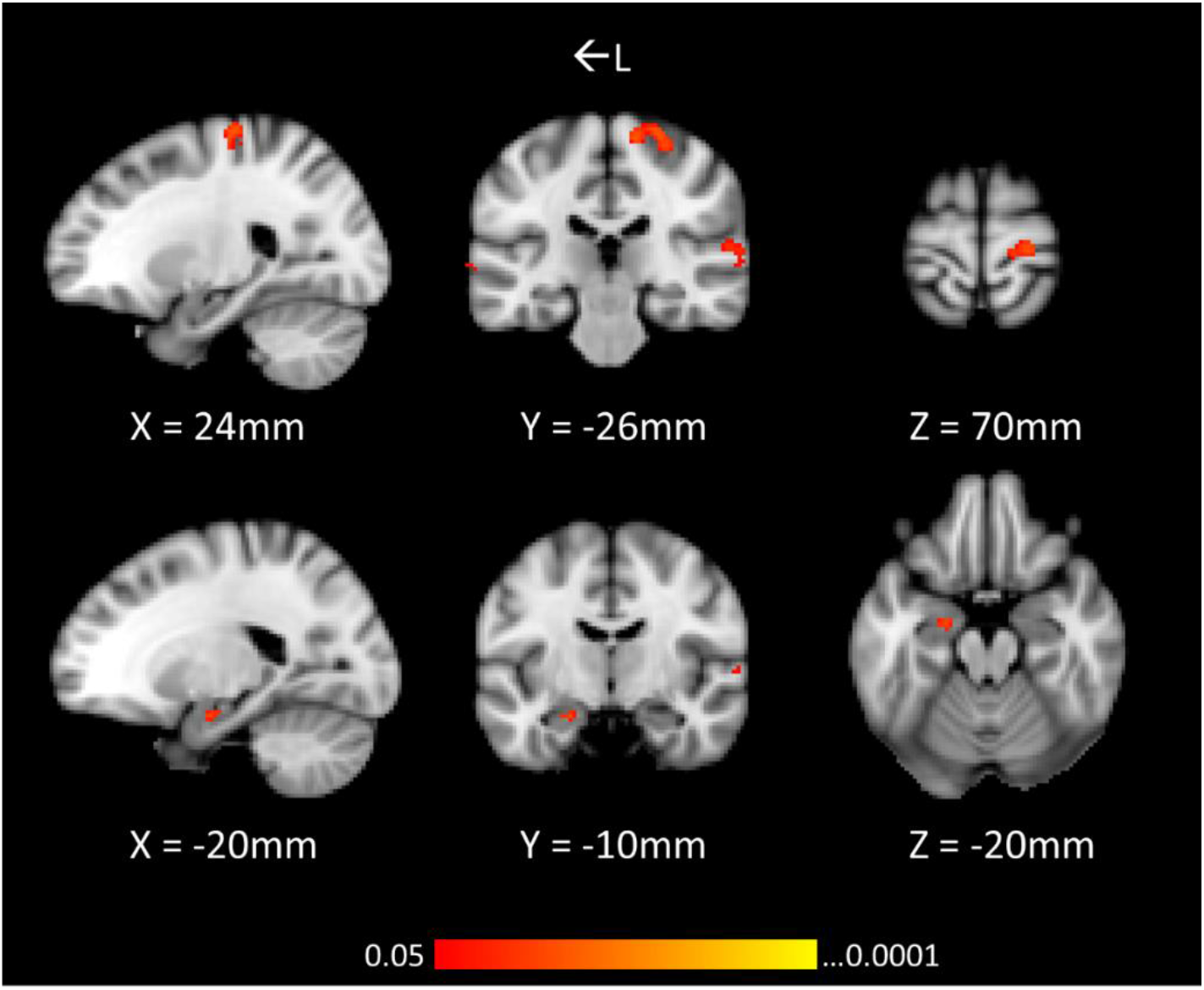
The (H1-Hsin) > (L1-Lsin) contrast yielded significant clusters in the right postcentral gyrus, right precentral gyrus, left middle temporal gyrus, right superior temporal gyrus, left hippocampus, and left amygdala. Top row coordinates are centered on right precentral gyrus, and bottom row coordinates are centered on left hippocampus.

Considering these activations in conjunction with our results from Experiment 1, these results suggest an interaction of value processing and statistical regularity, such that high-value first images (i.e., predictive images) in particular provoke deeper processing and greater attentional engagement than low-value predictive images. The greater activation in the IFG, left ACC, and LOC support our hypothesis that attention plays an important role in enhanced processing of the high-value first images.

Following up on these results, we examined how statistical regularities modulate responses, keeping value constant. We examined four contrasts: 1) H1 vs. Hsin, 2) H2 vs. Hsin, 3) L1 vs. Lsin, and 4) L2 vs. Lsin. We observed significant clusters for Lsin > L1 and Lsin> L2. The contrast of Lsin > L1 showed greater activation in middle temporal gyrus, hippocampus, amygdala, putamen, LOC, and other regions (Table 3 and Figure 8A), and the contrast of Lsin > L2 resulted in clusters in similar areas (Table 4 and Figure 8B). Comparisons between high-value paired images and high-value singletons did not yield any significant differences.

**Table 3.** Result of the contrast with Lsin>L1

| Anatomical Label | Hemisphere | Cluster size (voxel) | P value (TFCE) | Peak MNI, mm |
| --- | --- | --- | --- | --- |
| **Low-value singleton ($L_{sin}$) > Low-value first image ($L1$)** |  |  |  |  |
| Middle Temporal Gyrus, Superior Temporal Gyrus, Caudate, Parahippocampal Gyrus, Temporal Fusiform Cortex, Hippocampus, Amygdala, Cingulate Gyrus, Thalamus, Putamen, Postcentral Gyrus, Planum Temporale, Precentral Gyrus, Lateral Occipital Cortex, Ventricle, Right cerebellum, Accumbens | Both (unless stated specifically) | 44022 | 0.009 | 10, -34, 58 |
| Precuneous Cortex | Left | 146 | 0.036 | -22, -48, 26 |
| Lingual Gyrus | Left | 32 | 0.046 | -6, -70, 2 |
| Cerebellum | Left | 32 | 0.046 | -4, -46, -12 |
| Intracalcarine Cortex ,Precuneous Cortex | Right | 8 | 0.05 | 6, -66, 12 |

**Figure 8.**
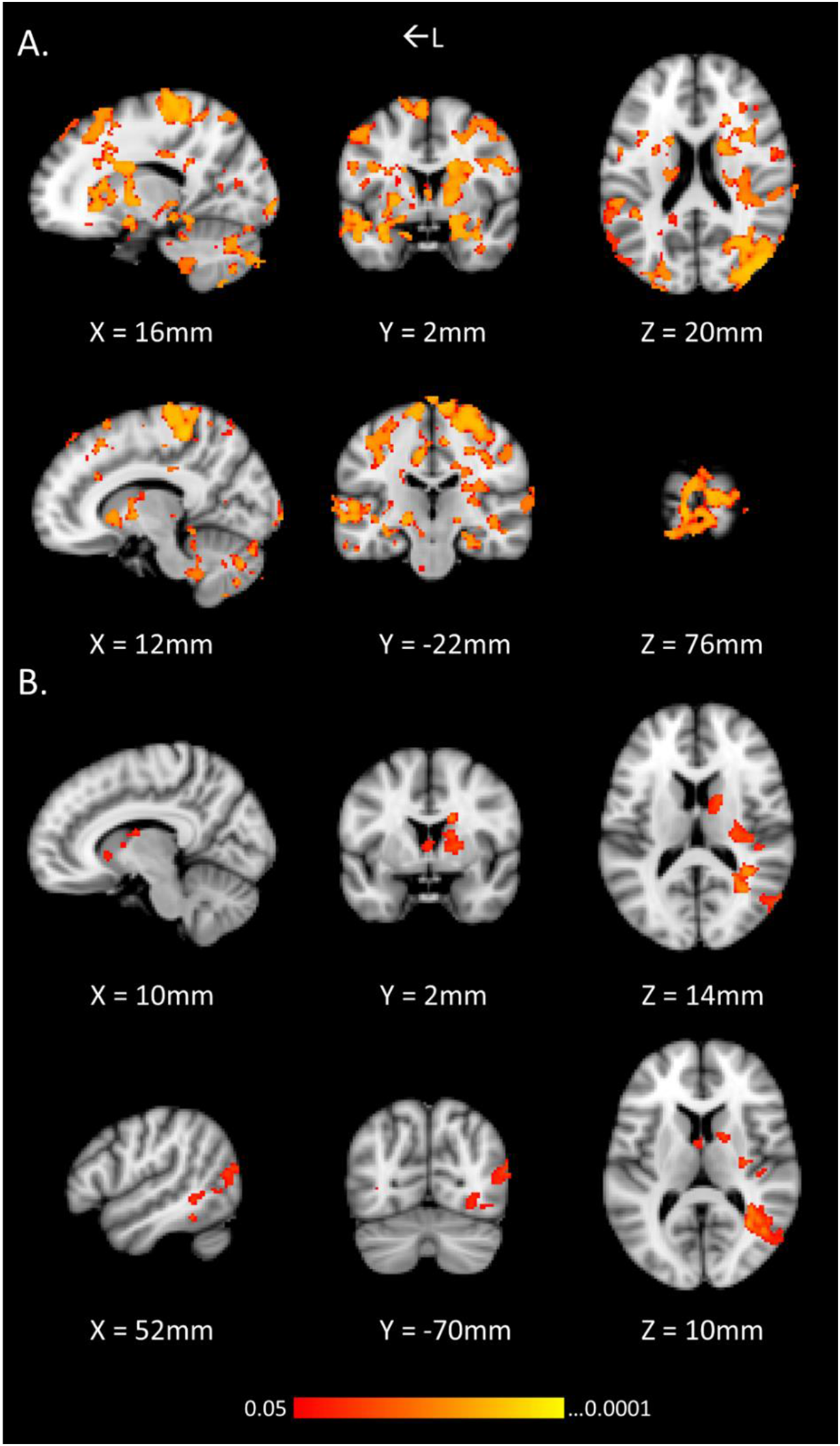
A) The contrast of Lsin > L1 showed greater activation in middle temporal gyrus, hippocampus, amygdala, putamen, and LOC. B) The contrast of Lsin > L2 also showed significant activations in middle temporal gyrus, hippocampus, inferior temporal gyrus, amygdala, putamen, and LOC. From top to bottom row, coordinates are centered on right caudate, right precentral, right caudate, and right LOC.

**Table 4.** Result of the contrast with Lsin>L2

| **Anatomical Label** | **Hemisphere** | **Cluster size**<br>**(voxel)** | **P value**<br>**(TFCE)** | **Peak MNI,**<br>**mm** |
| --- | --- | --- | --- | --- |
| **Low-value singleton (Lsin) > Low-value second image (L2)** |  |  |  |  |
| Middle Temporal Gyrus,<br>Superior Temporal Gyrus,<br>Precuneous Cortex,<br>Supramarginal Gyrus | Right | 1903 | 0.02 | 22, -52, 30 |
| Caudate, Pallidum, Putamen | Right | 670 | 0.021 | 16, 4, 26 |
| Ventricle | Both | 145 | 0.041 | 0, 4, 6 |
| Precuneous Cortex | Left | 63 | 0.038 | -24, -52, 26 |
| Orbitofrontal Cortex, Frontal<br>Pole, Caudate, Putamen | Right | 52 | 0.042 | 22, 32, -6 |
| Temporal Occipital Fusiform<br>Cortex | Left | 48 | 0.048 | -40, -44, -10 |
| Planum Polare | Right | 24 | 0.044 | 42, -24, -4 |
| Inferior Temporal Gyrus, Lateral<br>Occipital Cortex | Left | 6 | 0.049 | -48, -66, -14 |

We were not able to find any significant differences with contrasts scrutinizing the first images of paired images > singletons (see Turk-Browne et al., 2010). With our design, however, paired images contained not only statistical structure but also reward information, and the interaction between these two variables may drive a different pattern of results. Rather, we found that low-value singletons showed greater activity than low-value predictive (L1) images in areas recognized for playing a role in processing reward information (e.g., caudate, putamen, hippocampus). These results suggest that our (H1-L1)>(Hsin-Lsin) interaction may have been driven predominantly by differences in the way that L1 images are processed compared to low-value images that are non-predictive.

#### Passive viewing phase

During the passive viewing phase, participants were not required to perform any task other than to focus on each image as it goes by. We were interested in seeing whether any reward/structure related findings from the risky choice task phase would extend into other contexts (i.e., a context where participants are no longer making a choice or actively earning reward). However, we were unable to find similar patterns of activity with contrasts we ran with the risky choice task. We suspect that failure to observe patterns of activity similar to that found for the learning phase is possibly due to a lack of power, from only having time to collect data from a single run of the passive viewing task for most participants. This will be addressed further in the general discussion section.

#### Behavior data

We analyzed participants’ choices (i.e., yes or no) for the risky choice task for each time presentation (1st to 6th) collapsed over runs. As shown in Figure 9, a two-way repeated measures ANOVA (value of image x number of presentation) on risky choice proportion (i.e., choosing yes) showed a significant main effect of value, F (1, 28) = 51.21, p < .001, ηp^2^; = .647, and a trend (but not significant) of main effect of the number of presentation, F (1, 28) = 2.22, p = .055, ηp² = .074. A significant interaction between value of image (high or low) and the number of presentation (1st to 6th) was found, F (5, 140) = 29.46, p < .001, ηp² =.513. Proportion of making a risky choice was equally high for both high-value and low-value images at the first presentation, but across the second to sixth presentation, the proportion of making a risky choice on high-value images gradually increased, and the opposite was observed with low-value images.

**Figure 9.**
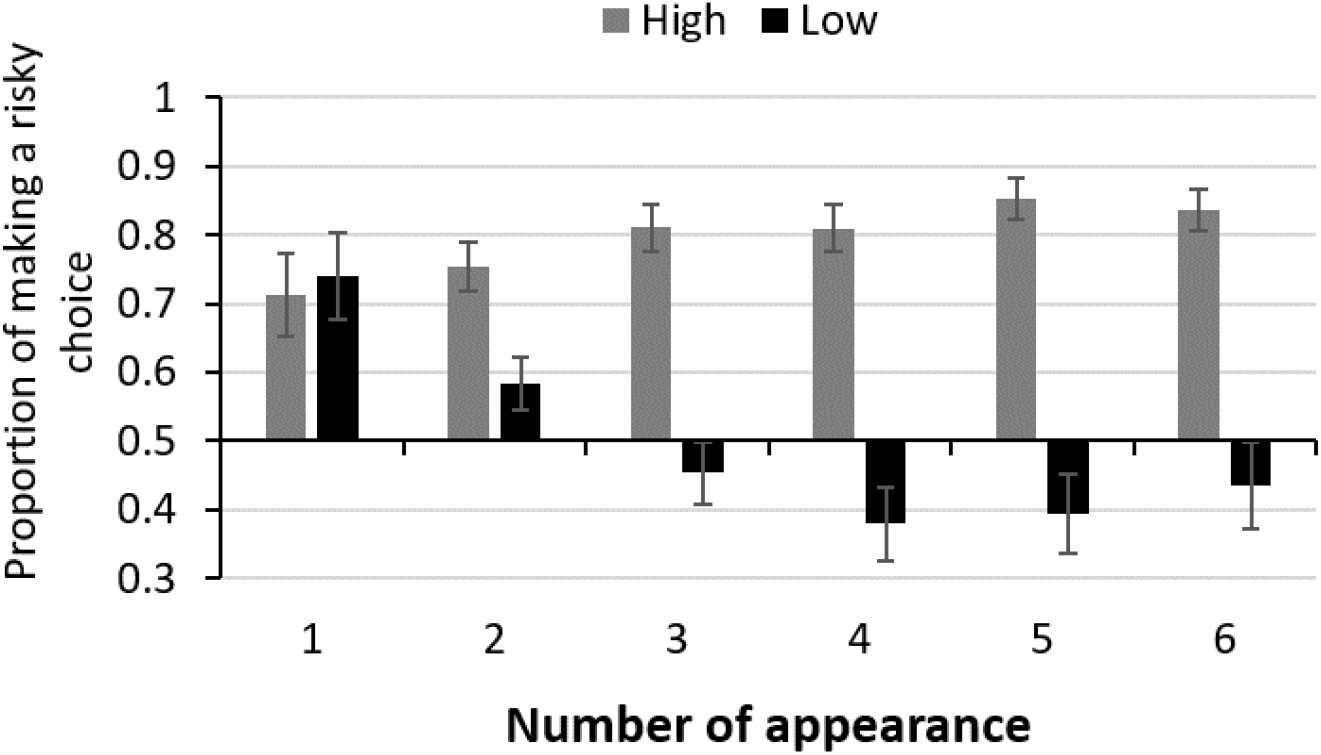
Proportion of making a risky choice by the number of presentations, split by value.

Participants completed two memory tasks after scanning: the recognition test and the reward memory test. As described earlier, there are several differences in design between Experiment 1 and Experiment 2, and based on these differences, the replication of behavior data was not guaranteed. In addition, we added the passive viewing task (one run: 21 participants; three runs: eight participants) after the learning phase, and this task might drive a different pattern of results.

For the recognition test, a one-sample t-test of recognition accuracy against chance (50%) yielded significant learning for all pair conditions; High-High: t(28)=3.3, p=.002, d=.62, High-Low: t(28)=3.26, p=.002, d=.6; Low-High: t(28)=3.1, p=.004, d=.57; Low-Low: t(28)=2.46, p=.02, d=.45 (Fig. 10). A 2 (value of first image, high or low) x 2 (value of second image, high or low) repeated measures ANOVA did not show any significant main effects nor an interaction (all p>.5; Fig. 10). In addition, a 2 (value of first image, high or low) × 2 (value of second image, high or low) repeated measures ANOVA did not reveal any main effects nor an interaction of foil type (i.e., foil pairs of High-High, High-Low, Low-High, and Low-Low conditions; F <1). Although the behavioral results of Experiment 1 did not replicate, this is likely due to design differences. This will be discussed further in the general discussion section.

**Figure 10.**
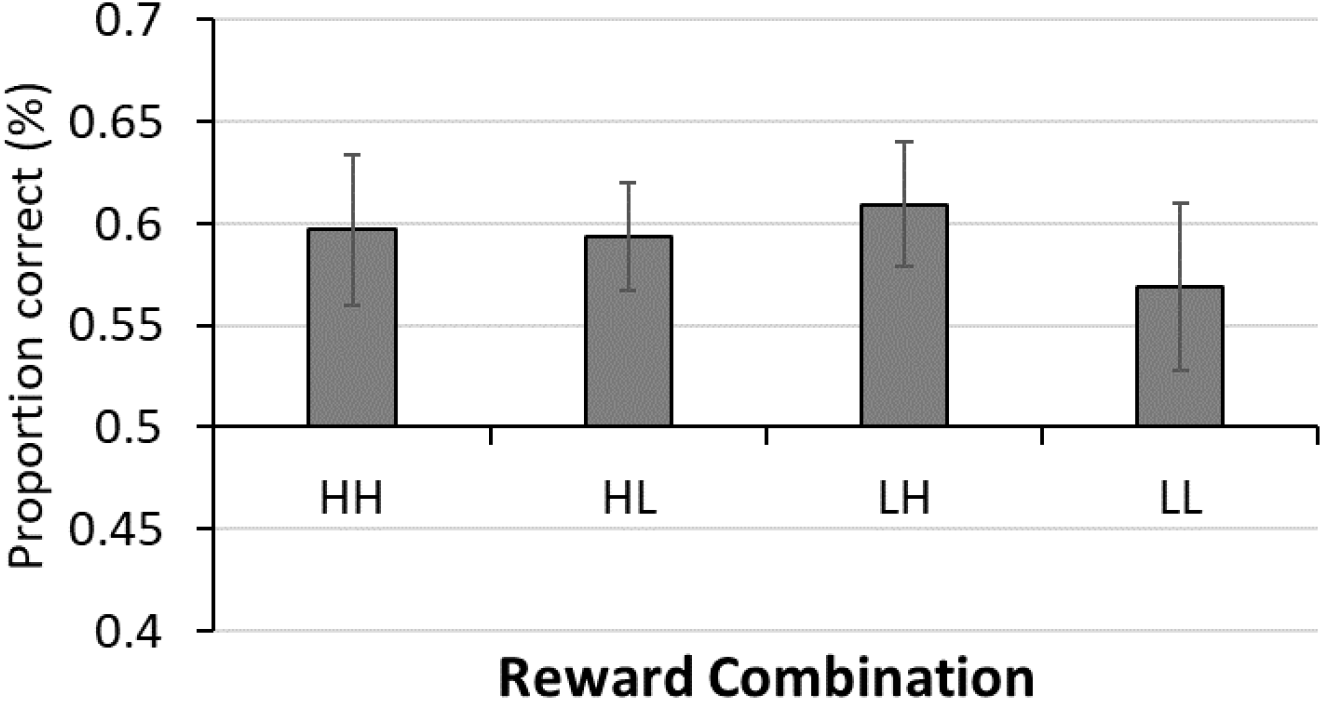
Accuracy at choosing target pairs over foil pair in four reward variations.

In the last reward memory phase, the mean proportion correct was 0.79 (SD: 0.13, t(28)=11.75, p < .001, d=2.18; one-sample t-test against chance, 50%). When we divided the results into the image type (the first, second images for pairs and singletons) and the reward type (high and low images), a repeated measures ANOVA did not show any significant main effect nor interaction (all p>.2; Fig. 11).

**Figure 11.**
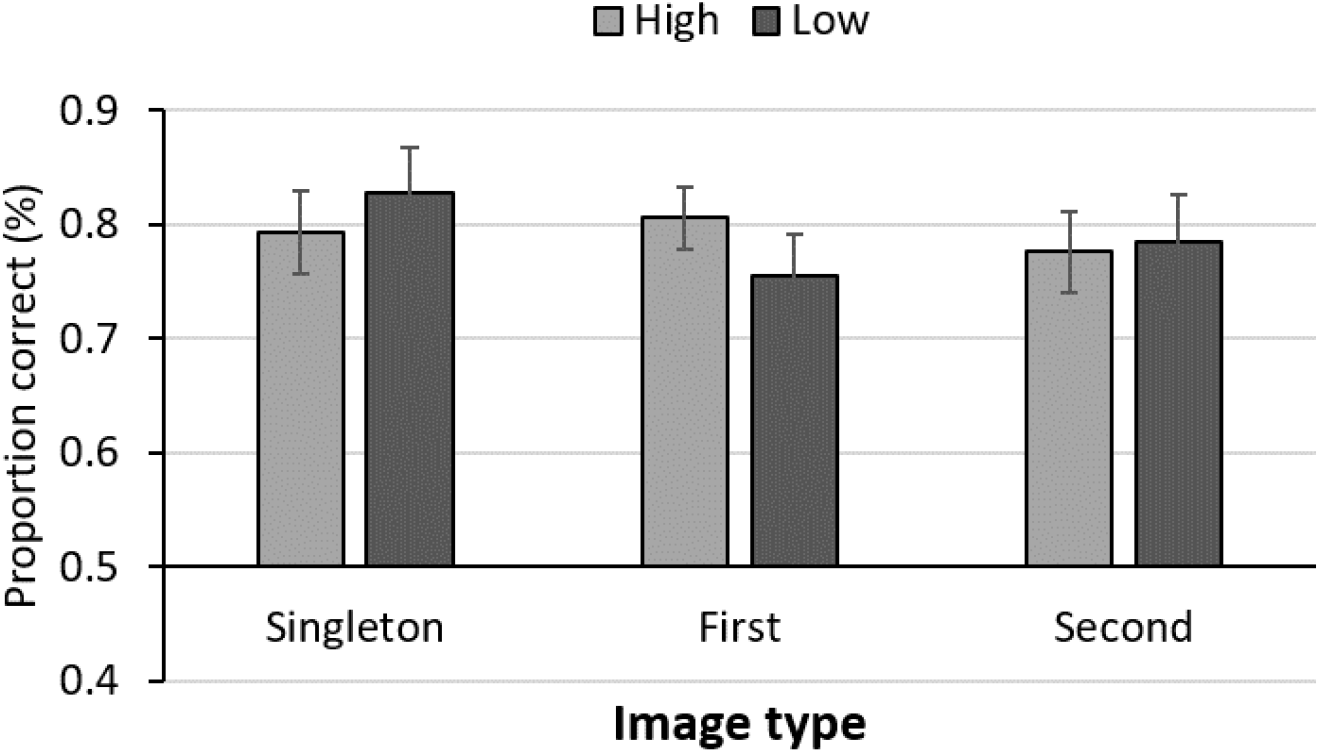
Accuracy at choosing reward value (high or low).

## DISCUSSION

In Experiment 2, we measured brain responses to visual images that were associated with both varying levels of reward and statistical contingencies. We found that the high-value first image (i.e., H1) led to greater activity in areas including IFG, left ACC, LOC, fusiform gyrus, orbitofrontal cortex (OFC), accumbens, precuneous cortex, parahippocampal gyrus, middle temporal gyrus, amygdala, hippocampus, and putamen as compared to the low-value first image (i.e., L1). These findings suggest that H1, in comparison to L1, led to greater attentional engagement (Beck & Vickery, 2020; Murray & Wojciulik, 2004; Stokes et al., 2009), and may enhance associative learning thusly. The contrasts of (H1-Hsin) > (L1-Lsin) yielded greater activations in the precentral gyrus, middle temporal gyrus, hippocampus, and amygdala, which supports the possibility that the differences between the high-value first image and the low-value first image are not driven solely by the value difference, but by an interaction of predictiveness and value. As no difference was found between the high-value singletons and the low-value singletons, the greater activation in precentral gyrus in this interaction also supports our hypothesis that the first image of the pair that was associated with high value received the attentional priority in comparison to that was associated with the low value (Fockert et al., 2004).

For contrasts comparing first images and singletons (e.g., H1>Hsin and L1>Lsin), we did not replicate the findings of Turk-Browne et al. (2010). We speculate that embedded reward information possibly altered learning in such a way that made it unique versus when VSL occurs in the absence of reward. The cover task was also quite different, which might contribute to differences in how VSL manifests. In addition, we found that low-value predictive images (i.e., L1) provoked less activity than non-predictive low-value singletons. In contrast, there was no difference in high-value comparisons between paired images and singletons. This suggests that the predictive nature of a stimulus may specifically down-regulate responses to low-value images, and thus that attention was less guided/prioritized to the low-value first image than the high-value first image.

## GENERAL DISCUSSION

Across two experiments, we provided behavioral and neural evidence that reward may alter visual statistical learning. In Experiment 1, better recognition of pairs when the first image of a pair was associated with high-value was observed, and this effect was especially pronounced for High-High pairs. We hypothesized that selective attention may play an important role in this finding, such that the first image of the pair that was associated with high value may receive the attentional priority in comparison to that was associated with the low value attention. Neural evidence supports our hypothesis, such that when the first image of a pair was associated with high-value, in comparison to the first image being associated with low-value (i.e., H1 > L1), greater BOLD response was observed in LOC and frontal and parietal areas, such as inferior frontal gyrus, precentral gyrus, and anterior cingulate gyrus, all regions whose activity is known to scale with attentional processing (Beck & Vickery, 2020; Corbetta & Shulman, 2002; Fockert et al., 2004). We did not observe similar differences with comparisons of H2 vs. L2, or Hsin vs. Lsin, which implies that not all high-value images led to attentional prioritization compared to low-value images. Rather, a combination of predictiveness and reward value was crucial in provoking this response. In VSL, the predictive image plays an important role, provoking anticipatory responses (Turk-Browne et al., 2010). Our findings suggest that VSL occurs differentially as a function of the magnitude of reward associated with the first image. In other words, when different amounts of reward are embedded in visual regularity, the rewards may interact with VSL in a way that it only impacts on the first position of the structured sequence, and selective attention may play an important role in formatting this interaction.

We additionally predicted that if greater attentional engagement occurs in the high value first image than the low value first image, it may yield greater activations in brain areas that have been known to play an essential role in associative learning (e.g., hippocampus, precuneous cortex, parahippocampal gyrus; Turk-Browne et al., 2010). Our results showed that in addition to the above areas, brain regions that have been known to play an important role in value processing (e.g., OFC, accumbens, and caudate; Baliki et al., 2013; Kringelbach & Rolls, 2004) showed greater activations in the high value image than the low value first image. Again, these findings are not driven solely by the value difference (i.e. high value vs. low value), but rather the interaction between statistically structured information and reward. With additional analyses, we showed that this value difference was specific to predictive items, suggesting that predictiveness and value interact in determining neural responses to images.

In Experiment 2, the processing of reward information elicited different patterns with behavior and neural approaches. Our behavioral results of learning of reward associations (i.e., the reward memory test) showed no difference in reward memory as a function of the structured information, which means that predictive structure (i.e., the first position of a pair vs. the second position of a pair vs. singleton) did not impact the recognition of reward information. The result of the reward memory test in Experiment 1 did not show any difference in reward memory of high or low images as a function of its position in a pair as well. However, our neural evidence reveals that reward-related responses were differentiated based on which structural position the reward was embedded in. We speculate that compare to the neural method, differences in the binding of reward based on stimulus-stimulus predictiveness may be too subtle for behavioral methods to uncover. The recognition task (i.e., two-alternative forced choice task) prior to the reward memory test might cause the subtler effect due to the presentations of foil pairs, which might interfere the reward memory as a function of the structured information.

Different results of the recognition phase between Experiment 1 and 2 may also be driven by design differences. For example, in Experiment 2, the gradual introduction of new pairs throughout the experiment might have cued learning and led to more generic learning effects. In addition, the timing of the risky choice task was different due to jittered intervals, and singletons were newly used in Experiment 2 as well. We presented singletons in between pairs during the learning phase to compare neural responses between paired images and singletons, but the inclusion of singletons might result in different patterns of behavior as compared to presenting only pairs, like in Experiment 1. Among these different experimental settings between Experiment 1 and 2, we suspect that the passive viewing task, where all pairs appeared one to three times across all participants, may play a critical role in yielding the different patterns. In this phase, participants were not required to make any choice, which means they did not have to process information related to reward variation. Hence, there is a possibility that the overall recognition rate may be increased across the board and eliminate the differences between reward variations.

To our knowledge, this is the first work to provide evidence of behavioral (Experiment 1) and neural responses (Experiment 2) being modulated by the interaction of reward and VSL. As mentioned above, Rogers et al. (2016) first explored the interaction between reward and VSL, but the reward variations (i.e., no-, low-, or high-reward) did not affect the learning of regularities. In our work, by using a risky choice task, we enhanced participants’ engagement to the task and value, and were able to observe an effect of reward on VSL. This implies that robust engagement with value information may be necessary to induce interactions with the learning of visual regularities. In conjunction with other recent results highlighting the importance of task during exposure shaping VSL (Vickery et al., 2018), the current study highlights the need to carefully consider context during exposure to regularities, and how those contexts shape incidental learning.

With respect to the passive viewing task, we were unable to observe a similar pattern of activity as that found in the risky choice task phase. We suspect that this failure is possibly due to a lack of power, due to our only having time to collect data from a single run of the passive viewing task for most participants. Another possible explanation for lack of such a finding is that the effect of reward in VSL may only arise within the context of tasks that draw attention to value, like our risky choice task. Therefore, simply viewing the sequence of images may not yield that same neural responses as actively making a risky choice on each image. Finally, because we introduced new sets of images in each learning run, it is possible that persisting VSL effects were variable across early vs. late, thus complicating detection of neural differences in our paradigm. Further studies of how the interaction between reward and VSL may affect the later representation of memory even outside of a reward-related context may be needed.

Another possible limitation in this study relates to the gambling task. There is an optimal response on high-reward trials but no optimal response for low-reward trials. In other words, the expected value for a yes response is higher than for a no response with high reward images, but the expected value was equal for yes and no responses with low reward images. However, the purpose of the gambling task is to enhance participants’ motivation to learn the value and to draw attention explicitly to reward during exposure. We, first, did not observe equal levels of choosing yes or no with low reward images even though the expected value is the same; rather, participants tend not to choose to gamble with low reward images throughout the task. Secondly, even if optimality may determine the different degrees of value learning, our neural results demonstrate that not only the higher value but also the order in the sequence matters, such that only the early items in a temporal sequence and value interact in determining neural responses to images.

The set of present studies provide evidence that VSL is modulated by reward. When a high reward is embedded in the first location of a statistically structured pair, it aids learning: a result we found support for in neural evidence. Several brain areas that reflect attentional capture, reward processing, associative learning, and the intermixed effect among them support the notion that reward contingencies affect VSL. These findings highlight the fact that reward may play a role similar to selective attention in VSL, such that the more the image can guide attentional resources, the better it can convey the reward information, and ultimately, facilitate visual statistical learning.

## Footnotes

1 Due to time constraints, twenty-one participants performed one run of passive viewing, and eight participants performed three runs of passive viewing.

2 However, when clusters were defined using a family-wise error (FWE) correction following a Z> 2.8 threshold (p < .005), based on Gaussian Random Field theory, we found greater activations in right LOC, with cluster size (voxels) as 443, Z= 3.776, and peak MNI (mm) on 42, −84, −8 (x, y, z).

## Notes

Author Note This research was supported by National Sciences Foundation grants BCS 1558535 and OIA 1632849

### Competing Interest Statement

The authors have declared no competing interest.

### Summary of Updates

This version of the manuscript has been revised to clarify the behavior findings.

